# Field-deployable molecular diagnostic platform for arbovirus and *Wolbachia* detection in *Aedes aegypti*

**DOI:** 10.1101/2020.04.28.066514

**Authors:** Natalie N. Rutkowski, Yuemei Dong, George Dimopoulos

## Abstract

**Background:** Surveillance of mosquito infection status is critical for planning and deployment of proper mosquito control initiatives. Concurrently, *Wolbachia* is being widely used as a control method for arboviral transmission. Point-of-care (POC) detection assays are necessary for monitoring the infection prevalence and geographic range of viruses as well as *Wolbachia* in mosquito vector populations. We therefore assessed the novel qPCR bCUBE molecular diagnostic system as a tool for virus and *Wolbachia* detection.

**Results:** We developed a reliable, specific, and sensitive diagnostic assay for detecting Zika virus and dengue virus serotype 2 using the real-time qPCR platform bCUBE. With bCUBE-based qRT-PCR, both *Wolbachia* bacterium and virus RNA could be reliably detected in individually infected *Ae. aegypti* mosquitoes and in pools of 5, 10, or 15 mosquitoes.

**Conclusions:** The portable qPCR bCUBE diagnostic platform is capable of detecting Zika and dengue virus as well as *Wolbachia* in mosquitoes and therefore has potential as a practical field-deployable diagnostic test for vector-borne disease surveillance programs.

## Introduction

Arthropod-borne diseases threaten over two-thirds of the global population and are exhibiting an ongoing expansion of their geographic range and prevalence as a result of climate change, urbanization, and globalization (1–4). According to the World Health Organization, dengue virus infection has increased 30-fold over the past 50 years and now affects nearly 100 million people, primarily in the Americas and Asia (5, 6). Recent outbreaks of Zika virus have also led to widespread concern because of the virus’s ability to cause newborn malformations (7, 8).

Dengue and Zika viruses are members of the genus *Flavivirus* that are primarily transmitted by the *Aedes* mosquito (7, 9, 10). Other members of the *Flavivirus* genus are also recognized as vector-borne pathogens of public health significance, including West Nile virus (WNV), yellow fever virus (YFV), Japanese encephalitis virus (JEV), and Chikungunya virus (CHIKV). These viruses result in similar flu-like symptoms with the potential to progress to neuroinvasive outcomes. *Aedes* mosquitoes, with their aggressive blood-feeding behavior, have allowed for efficient human-mosquito transmission of these arboviruses (3, 11). The geographical presence of this mosquito vector has dramatically increased in the last few decades, leading to an expanded transmission of these arboviruses (12–14). The lack of vaccines and treatment against these arboviruses highlights the importance of mosquito control and surveillance strategies (15). Current and future mosquito-targeted control strategies are, and will be, having a significant epidemiological impact but also require mosquito and pathogen surveillance (15). Surveillance of geographical distribution of the vector mosquitoes and the pathogens they carry is an essential component of disease prevention and control.

*Wolbachia pipientis* is increasingly being used as a method for limiting arboviral transmission in dengue and Zika virus-endemic countries. This endosymbiotic bacterium is passed from mother to offspring and has been shown to suppress dengue and Zika virus transmission in *Aedes* mosquitoes (16, 17). Field-release trials require continual monitoring of *Wolbachia-infected Ae. aegypti* in mosquito populations (18–20). Real-time qPCR has been used as the primary method for detecting *Wolbachia* genetic material in mosquitoes (21).

Accurate, rapid, and cost-effective mosquito and pathogen surveillance is critical for monitoring infection prevalence and thereby mitigating transmission risk. Methods currently used for arboviral detection include viral culture, antibody detection, antigen detection, and RNA detection using quantitative real-time RT-PCR (qPCR). Currently, the qPCR method is the gold standard because of its use of specific molecular markers (22, 23). Many of these assays are laborious and laboratory-based and require expensive and bulky instruments, making them incompatible with low-resource regions (24). Recently, several novel, advanced point-of-care (POC) diagnostic measures have been developed for detecting mosquito-borne viruses in the field, including honey-baited nucleic acid preservation cards (25, 26), loop-mediated isothermal amplification (LAMP) (27), biosensors (28), and adaptations of near-infrared spectrometry techniques (29). However, these techniques have documented limitations, including cross-reactivity with other flaviviruses or requirements for training on specialized equipment, making the adoption of these new diagnostic tools difficult (30, 31). Although the global burden of emerging outbreaks of Zika and dengue is clearly recognized, there is a gap in resources for endemic countries and regions, which are consistently plagued by a lack of equipment and adequate resources to consistently monitor the prevalence and range of infected mosquitoes (5, 32, 33).

Disease surveillance and integrated vector control are essential for curbing disease transmission. Nucleic acid-based testing to detect viral RNA allows for specific and sensitive virus monitoring in mosquito surveillance programs (24). Several conventional and more recent real-time PCR-based assays have been established for mosquitoes and their vectored pathogens, and the ready availability of genome sequences for both vectors and pathogens can support the identification of additional PCR-compatible molecular markers (34–36).

In the present study, we evaluated a novel and portable real-time PCR platform, bCUBE (Hyris, Ltd), as a PCR-based arboviral detection method with potential for field-deployability. bCUBE makes possible the genetic testing of biological samples in any setting, at any time, with real-time access to results on its dedicated cloud-based software platform. This technology is a portable device, similar in size to a Bluetooth speaker, which is used for biological analysis in several fields, including agricultural pest control. The device can be operated from a laptop, tablet, or cellular device through an easy-to-use gateway and generates centralized data analysis immediately after a reaction. The device is capable of performing thermocycling reactions such as real-time PCR, as well as loop-mediated isothermal amplification (LAMP). The system can be calibrated to distinguish between positive and negative samples in a single reaction with predetermined conditions that have been established ahead of time in the laboratory. This feature allows the bCUBE to be operated by individuals lacking in depth training in qPCR assays and data analysis skills. We have now explored the use of bCUBE technology for detection of both dengue and Zika virus in *Aedes aegypti*, optimizing and standardizing the sample preparation method to be used with a commercially available one-step qRT-PCR assay kit. Finally, we have developed a bCUBE-compatible qPCR diagnostic assay for the surveillance of arboviral pathogens and *Wolbachia* in *Aedes* mosquitoes.

## Methods

### Ethical Statement

This study was carried out in strict accordance with the recommendations in the Guide for the Care and Use of Laboratory Animals of the National Institutes of Health (NIH). Mice were used according to an animal protocol (permit # MO15H144) approved by the Animal Care and Use Committee of the Johns Hopkins University and were used as the blood source for the maintenance of the mosquito colonies. Commercial anonymous human blood (InterState Blood Bank) was used for Zika and dengue virus infection assays in mosquitoes, and informed consent was therefore not applicable.

### Mosquito rearing and mosquito infections

*Ae. aegypti* Liverpool strain LVPB12 mosquitoes were maintained on 10% sucrose solution under standard insectary conditions at 27±0.5°C and 75-80% humidity with a 12-h light/dark cycle. Mosquitoes were reared using a standard rearing protocol, and colonies were maintained on Swiss Webster mice (Charles River Laboratories). DENV2 and ZIKV infection of mosquitoes was carried out as previously described (37). Virus supplemented blood was fed to mosquitoes using an artificial membrane glass feeder, and infected mosquitoes were double-caged and incubated in a reach-in incubator under conditions similar to those in the standard insectary chamber described above. *Wolbachia w*AlbB-infected *Ae. aegypti* (WB1) mosquitoes were reared and maintained under the standard insectary conditions described above and in (38).

### Cell culture and virus propagation

*Aedes albopictus* C6/36 cells (ATCC CRL-1660) were cultured, and viral stocks were prepared as previously described in (39). In brief, C6/36 cells were cultured in MEM medium (Gibco) supplemented with 10% heat-inactivated fetal bovine serum (FBS), 1% penicillin-streptomycin, and 1% non-essential amino acids and maintained in a tissue culture incubator at 32°C and 5% CO_2_. Baby hamster kidney strain 21 (BHK-21, ATCC CCL-10) cells were maintained at 37°C and 5% CO_2_ in the DMEM medium (Gibco) supplemented with 10% fetal bovine serum (FBS), 1% penicillin-streptomycin, and 5μg/ml Plasmocin™. DENV serotype 2 New Guinea C strain (DENV2) and ZIKV strain FSS 13025 (ZIKV) were used in the indicated experiments. For viral stock preparation, C6/36 cells grown to 80% confluence were infected with ZIKV and DENV2 at a multiplicity of infection (MOI) of 10 and incubated at 32°C and 5% CO_2_ for 5 days (DENV2) and 6 days (ZIKV). Virus was harvested by three freeze-thaw cycles using dry ice and a water bath (37°C), then centrifuged at 2,000 rpm for 10 min at 4°C. The supernatant from this cell lysis was mixed with the original cell culture supernatant to yield the final viral stock. Viral stocks were aliquotted and stored at −80°C for long-term storage. Viral stock titration was done by plaque assay as described below.

### Primer design for real-time quantitative RT-PCR (qPCR)

The ZIKV envelope (E) protein was chosen as the target for ZIKV detection, and primers were developed based on previously established sequences (40). Dengue virus serotype 2 (DENV2) qPCR was performed using previously developed protocols that employ primers detecting serotype 2 in the 3’UTR region (41) (Table 1). The primers were modified and optimized for bCUBE qPCR (Table 1). Primer sequences were assessed with NCBI Blast to search for potential cross-reactivity against other viruses, bacteria, and vectors. Previously established primers for *w*AlbB detection were used for *Wolbachia-infected Ae. aegypti* (42). The sequences of the primers are summarized in Table 1.

**Table 1.**
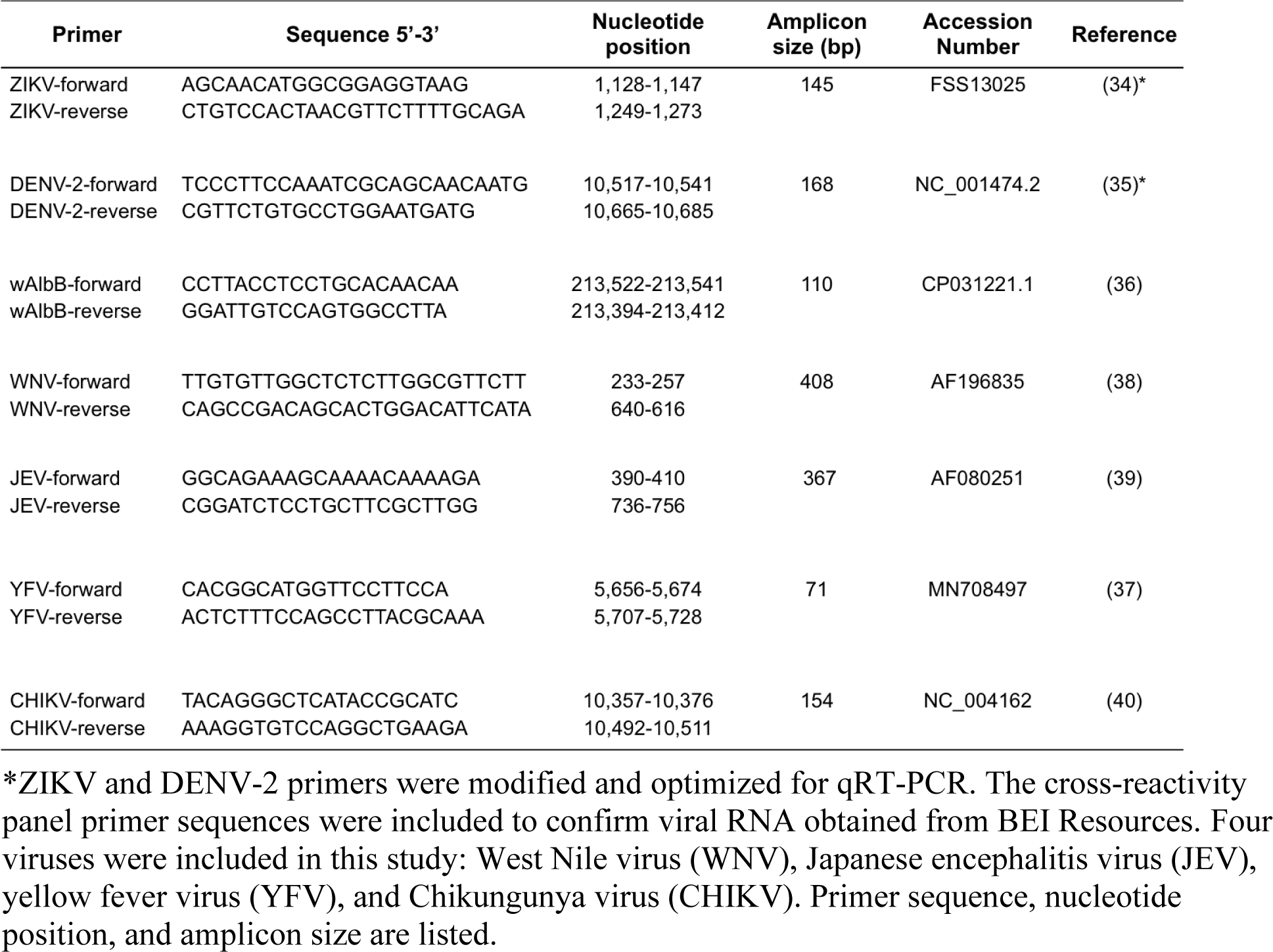
Primer pairs for viral detection and cross-reactivity panel for real-time qPCR using the bCUBE.

### Cross-reactivity panel of mosquito-borne viruses

Viral nucleic acid sequences included in the cross-reactivity panel were CHIKV (H20235 ST MARTIN 2013), JEV (India R53567), WNV (CO 1862), and YFV (17D), obtained from BEI Resources. A NanoDrop Spectrophotometer (ThermoFisher Scientific) was used to measure the RNA concentration of these viral samples. Previously established primers were used, as listed in Table 1, for qPCR to confirm the presence of corresponding viral RNA in the samples (43–46).

### Total RNA preparation for DENV2- and ZIKV-infected whole mosquitoes or tissues

Total RNA extraction was performed using a squash buffer (10 mM Tris base, 1 mM EDTA, 50 mM NaCl) supplemented with 1:8 part Proteinase K (Qiagen, 15mg/ml). Mosquito abdomens with the midguts and heads with thoraces were individually collected in 50 μl of squash buffer at 7 and 14 days, respectively, after an infectious blood meal and stored at −80°C until extracted. Proteinase K was added at 1/8 volume to each sample to give a final concentration of 15 mg/ml and homogenized with a cordless Pellet Pestle Motor (Kontes) for 40-60 s. Samples were incubated at 57°C for 5 min, followed by 95°C for 5 min for enzyme deactivation. The supernatant from this crude RNA extraction was used immediately for qPCR or stored at −80°C until use.

### gDNA preparation of *Wolbachia*-infected mosquitoes

Total gDNA from *Wolbachia-infected* total mosquitoes or tissues was prepared as described above (47). These crude gDNA extractions were immediately used for qPCR or stored at −80°C until use.

### Laboratory standard real-time PCR with Applied Biosystems equipment

A StepOnePlus Real-Time PCR system (Applied Biosystems) was used for comparative analysis. The mosquito housekeeping gene ribosomal protein S17 (*rps17)* was amplified in *Aedes aegypti* using the two different PCR machines (Applied Biosystems and bCUBE) with the same real-time PCR settings. SYBR Green PCR Master Mix (Applied Biosystems) was used according to manufacturer’s instructions. Mosquitoes were collected individually in 50 μl of squash buffer, and crude gDNA was extracted. Identical samples were applied to both the bCUBE- and StepOnePlus-based qPCR systems for comparative evaluation.

### Hyris bCUBE real-time RT-PCR

The portable qPCR machine Hyris bCUBE 2.0 thermocycler (Hyris, London, UK) and the 16-well cartridges were kindly provided by Hyris Inc. The Hyris data analysis platform was used for this study. GoTaq 1-Step RT-qPCR kit (Promega) was used at a final volume of 20 μl with: 0.4 μl of kit reverse transcriptase enzyme, 10 μl of RT-PCR buffer, 0.3 μl of 10 μM of forward and reverse primer, 9 μl of DNA/RNA free water, and 1 μl of crude RNA or gDNA sample. Each sample was performed in technical duplicate. Each cartridge run included one negative and one positive control. The following thermocycling settings were used: reverse transcription for 15 min at 37°C, heat-inactivation of reverse transcriptase at 95°C for 10 min, followed by 30 PCR cycles at 95°C for 10 s, 60°C for 30 s, and 72°C for 30 s. Melting curve analysis was performed at the end by cooling to 60°C, followed by heating to 95°C at 0.05°C/s. Automatic data analysis was generated with the Hyris data analysis platform. Two technical replicates were performed on the platform for each biological replicate. Three biological replicates were included for each assay.

### Absolute quantification of viral copy numbers through qRT-PCR

*In vitro-*transcribed RNA of the E gene of ZIKV and the 3’-UTR region of DENV2 were used for absolute quantification and cloned into a TOPO-TA vector. Total RNA was extracted using TRIzol reagent (ThermoFisher Scientific). RNA (2 μg) was used for cDNA synthesis with M-MLV Reverse Transcriptase (Promega). Conventional PCR was used with the primer (Table 1) to amplify a DNA fragment of 168-bp for DENV2 and 145-bp for ZIKV. PCR was followed by PCR purification using a QIAquick PCR Purification kit and protocol (Qiagen). About 50 ng of cleaned PCR product of either DENV2 or ZIKV was separately cloned into a TOPO-TA PCR cloning vector (ThermoFisher Scientific) according to the manufacturer’s instructions. The ligation product was transformed with TOP10-competent cells (ThermoFisher Scientific), and positive clones grown on LB selection agar plates were screened through colony-PCR, followed by plasmid mini-prep (Qiagen) and sequencing confirmation. The plasmids of the final confirmed-positive clones were purified using the Maxiprep kit (Qiagen) according to the manufacturer’s instructions. Plasmid DNA was used to prepare absolute standards. A HiScribe T7 High Yield RNA Synthesis Kit (NEB) was used for RNA synthesis, and the concentration of the RNA was checked with a NanoDrop Spectrophotometer (ThermoFisher Scientific). Molecular weights were converted to copy numbers using the New England BioLabs Calculator (https://nebiocalculator.neb.com/#!/ssrnaamt). The concentration of RNA was adjusted to 10^10^ copies/μl and serially diluted 10 times for the standard curve on the bCUBE. The standard curve was generated using GoTaq 1-Step RT-qPCR (Promega).

### Viral titration by plaque assay

The titers of ZIKV and DENV2 in the original viral stocks and in the infected mosquitoes were determined by plaque assays in BHK-21 cells. Each experiment was performed with at least three biological replicates. Whole mosquito or mosquito tissue samples were collected at 7 and 14 days post-infectious blood meal in 150 μl of complete DMEM medium with glass beads. Tissue samples were homogenized with a Bullet Blender (Next Advance) and serially diluted with DMEM complete medium. One or two days before plaque assay, the BHK-21 cells were split at a 1:10 dilution and grown on 24-well plates to 80% confluency. Serially diluted mosquito or viral samples (100 μl each) were added to the BHK-21 cells, followed by incubation at room temperature for 15 min on a rocking shaker (VWR) and subsequent incubation at 37°C with 5% CO_2_ in a cell incubator (ThermoScientific) for another 45 min. The 24-well plates with infected BHK-21 cells were overlaid with 1 ml of 0.8% methylcellulose in complete DMEM medium with 2% FBS and incubated for 5 to 6 days in the cell culture incubator (37°C and 5% CO_2_). Plaques were fixed and developed with staining reagent (1% crystal violet in 1:1 methanol/acetone solution) at room temperature for approximately 30 min. Plates were rinsed with DI water and air-dried, and plaques were counted and multiplied by the dilution factor to calculate the plaque forming units (PFUs) per sample.

### DENV2 and ZIKV mosquito pools

Preliminary pooled sample experiments involved a total of 300 *Aedes aegypti:* 276 uninfected, 12 infected with DENV2 and 12 infected with ZIKV. First, 12 mosquitoes were infected with ZIKV and 12 with DENV2. Following confirmation of infection by RT-qPCR of individual infected mosquitoes, each was placed into a pool of uninfected mosquitoes. Four different pools were used to measure the sensitivity of the infection detection for pools of 5, 10, 15, and 20. The squash buffer volumes used were 250, 450, 700, and 950 μl, respectively, followed by the addition of 1:8 part Proteinase K.

### Statistical Analysis

Graphs were generated using GraphPad Prism Software version 8. The estimation of several diagnostic parameters for leg samples, including sensitivity, specificity, accuracy, and positive and negative predictive values was calculated using the web-based software MedCalc Diagnostic Test Evaluation Calculator.

## Results

### The reliability of the portable bCUBE qPCR machine is equal to that of a laboratory-standard qPCR system

The reliability of the novel bCUBE real-time PCR system was validated by comparing the system to a standard qPCR instrument (Applied Biosystems StepOnePlus, ABI). Our proof-of-principle studies used mosquito tissue homogenates (abdomen with the midgut, and head with the thorax) in squash buffer that were tested with the mosquito housekeeping gene (ribosomal protein S17, *rps*17). *Aedes aegypti* were dissected into crude compartments, including the abdomen and head plus thorax. Identical samples were processed using both qPCR machines, identical settings, and identical SYBR Green Master Mix and instructions (Applied Biosystems). As shown in Fig. 1, the housekeeping *Rps*17 gene was detected in all 24 samples by both machines. The cycle threshold (Ct) values for the *Rps*17 gene in *Ae. aegypti* were detected significantly earlier in the bCUBE system than in the laboratory standard procedure, with a mean difference of 2.40 Ct value (paired t-test, *p*-value < 0.0001). This result suggests that the bCUBE qPCR is reliable in detecting a mosquito gene from a single mosquito tissue using a crude sample-preparation method.

**Fig. 1.**
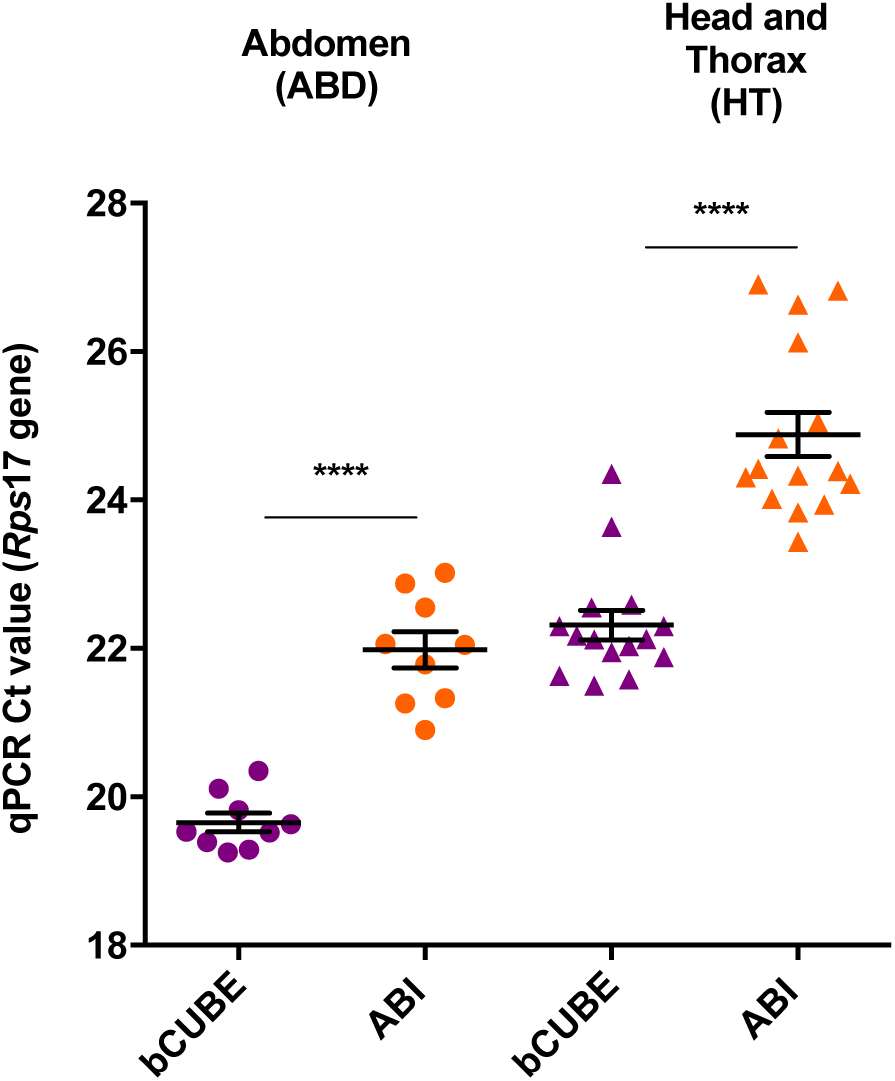
Comparison of *Ae. aegypti Rps*17 gene detection in abdomen (ABD) and head plus thorax (HT) samples using bCUBE and Applied Biosystems real-time PCR. *Ae. aegypti* abdomens and head plus thoraces were collected, and gDNA was extracted using the squash buffer method. The housekeeping gene *Rps*17 was amplified using the bCUBE and Applied Biosystems real-time PCR systems. Cycle threshold (Ct) values are plotted. Statistical significance was determined by Mann-Whitney test (****, p < 0.0001; Mann-Whitney).

### bCUBE-based one-step qRT-PCR is sufficiently sensitive for DENV2 and ZIKV detection

We performed sensitivity studies on the bCUBE-based qPCR system based on absolute quantification standard curve analysis. Cloned fragments of DENV2 and ZIKV were quantitated for RNA copy number and serially diluted to establish a qPCR standard curve. End-point limitations were set to < 30 for Zika virus, whereas dengue end-point limitations were set to < 29 based on standard curve (Supplementary Fig. 1). Correlation coefficient (R^2^) values were 0.99 for DENV2 and 0.98 for ZIKV. Based on the standard curve analyses, the limit of detection range was 10 viral RNA copy numbers for DENV2 and ZIKV. Amplification above these values was attributed to non-specific amplification.

### bCUBE-based qRT-PCR is specific for DENV2 and ZIKV detection

The specificity of the ZIKV and DENV2 assays was then determined using a cross-reactivity panel. To address challenges involving primer cross-reactivity with other arboviruses (46, 48) as well as the tendency toward false-positive amplification in negative samples, we tested the specificity of the assay against a panel of *Flaviviruses* and one *Alphavirus*. The frequently co-circulating arboviral RNA samples, obtained from BEI resources (Table 2) for this study, were: CHIKV (H20235 ST MARTIN 2013), JEV (India R53567), WNV (CO 1862), and YFV (17D). The ZIKV and DENV2 primer pairs showed a high degree of specificity for amplification of their respective virus RNAs (Table 2), since no amplification of other virus RNAs occurred.

**Table 2.** Evaluation of the specificity of bCUBE-based qRT-PCR for DENV2 and ZIKV detection. A cross-reactivity panel of frequently co-circulating viruses was used to evaluate the specificity of bCUBE-based qRT-PCR for DENV2 and ZIKV. Arboviral RNA samples were obtained from BEI Resources. The presence of all cross-reactivity panel RNA samples was quantified on NanoDrop for molecular weight and amplified on qRT-PCR with their own primer sequences.

| Family | Genus | Species | Strain | BEI No. | Result for ZIKV qPCR Assay | Result for DENV-2 qPCR Assay |
| --- | --- | --- | --- | --- | --- | --- |
| *Flaviviridae* | *Flavivirus* | Zika virus |  |  | + | - |
|  |  | Dengue virus serotype 2 |  |  | - | + |
|  |  | Japanese Encephalitis virus | India | NR-9592 | - | - |
|  |  | West Nile virus | CO 1862 | NR-50434 | - | - |
|  |  | Yellow Fever virus | 17D | NR-2869 | - | - |
| *Togaviridae* | *Alphavirus* | Chikungunya virus | St Martin 2013 | NR-50130 | - | - |

### bCUBE-based qRT-PCR is as accurate as plaque assay in detecting infectious viral RNA

PCR detection of non-infectious viral RNA is possible when viral RNA is not actively replicating (49). Therefore, the viral RNA copy number and viral infectious RNA load can differ in mosquitoes. We thus sought to evaluate the infection prevalence using bCUBE qRT-PCR and plaque assays as opposed to titer comparisons. Three biological replicates of *Ae. aegypti* were infected with ZIKV and DENV-2. Each replicate of infected mosquitoes was collected and separated into two halves to compare infectious prevalence as determined by plaque assay and bCUBE qPCR analysis. In each of the three replicates, no-significant differences were seen in infection prevalence values for both ZIKV and DENV2 between the two methods. These results suggest that bCUBE qRT-PCR analysis can accurately detect infectious ZIKV AND DENV2 (Fig. 2).

**Fig 2.**
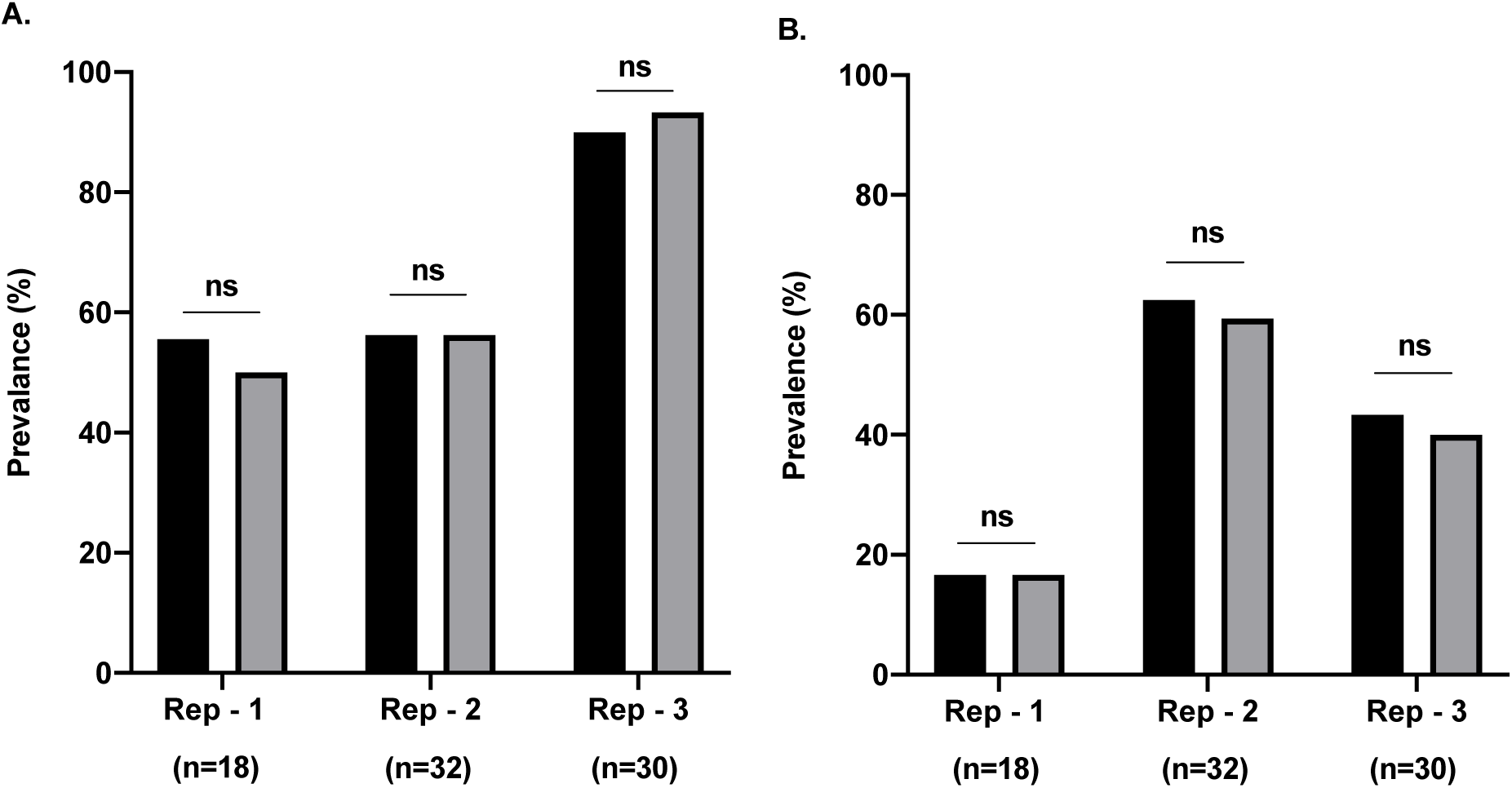
ZIKV AND DENV2 infection prevalence in *Ae. aegypti* as detected in the bCUBE versus plaque assay. *Ae. aegypti* were infected with Zika (A) and dengue (B) virus via an artificial blood meal. Each biological replicate was split into two groups and analyzed using the bCUBE assay (black bars), and the other half was used for the plaque assay (grey bars). No significant difference was detected between the plaque assay and bCUBE qPCR in terms of infectious prevalence (ns: not significant; Fisher’s exact test).

### ZIKV and DENV-2 RNA can be detected and quantified in individual mosquito samples by bCUBE assay

*Ae. aegypti* mosquitoes were experimentally infected with ZIKV (n=112) and DENV2 (n=112) through an artificial blood feeder containing anonymous human blood supplemented with virus, and samples from the mosquitoes were evaluated for infection status with the bCUBE platform at 7 and 14 days post-blood meal. Samples with Ct values < 30 were considered positive for ZIKV and < 29 for DENV2 infection. Each sample was run in duplicate to confirm infection. Both ZIKV and DENV2 viral RNA were detected from an individually infected mosquito using the squash buffer extraction method and the bCUBE platform. Viral RNA could be detected and quantitated in samples containing single abdomen with midguts and head with thoraces of infected mosquitoes (Fig. 3).

**Fig 3.**
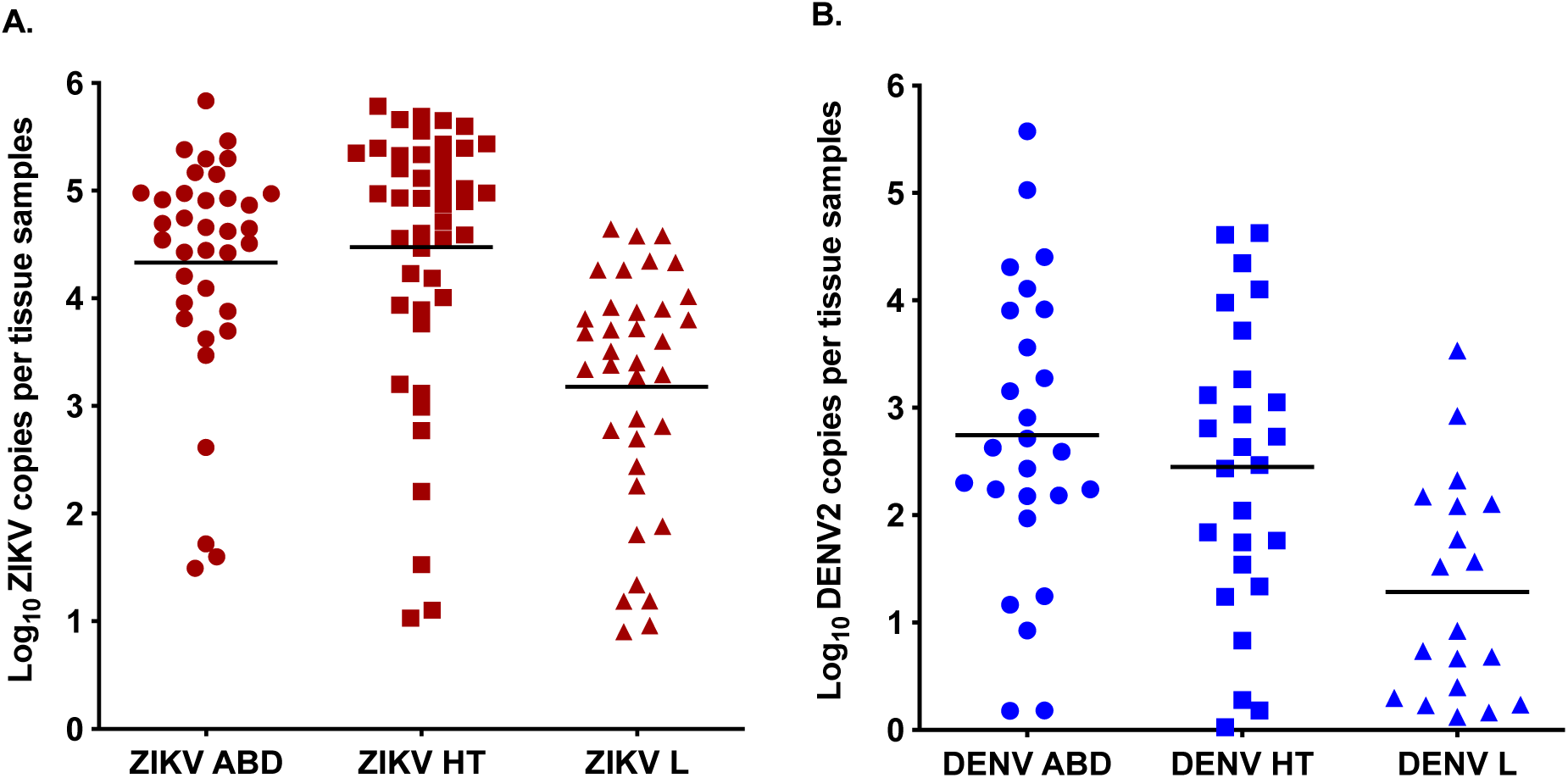
Viral RNA concentrations of individual *Ae. aegypti* tissues collected at various time points. Individual *Ae. aegypti* that were infected with Zika (A) and dengue virus (B) were collected at 7 or 14 days post-infectious blood meal. Abdomens with midgut samples (ABD) were collected on day 7 post-infectious blood meal, and head with thorax samples (HT) were collected on day 14. Corresponding leg (L) samples were also collected on day 14. Viral RNA extraction was performed using the squash method, followed by amplification using 1-step qPCR on the bCUBE. Uninfected mosquitoes were not included.

Next we investigated detection of virus in leg samples of *Ae. aegypti* (Fig. 3). Forty-six leg samples were collected from mosquitoes that were also analyzed for infection in the heads and thoraces to confirm infectious status. We found significant differences between the Ct values for the head with thorax samples versus the leg samples. For the leg samples, six false negative sample results were obtained for DENV2, and two false negative results were obtained for ZIKV. One false positive result was detected for ZIKV (Fig. 4). These results indicate that analysis of infection using leg samples is not reliable. No amplification occurred in uninfected *Aedes aegypti* used as a negative control. Because of the false negative results, the DENV2 sensitivity of leg samples was 76% (95% CI, 54.87% to 90.64%) and the diagnostic specificity of leg samples was 100% (95% CI, 83.89% to 100.00%). The sensitivity for ZIKV of leg samples was 94.59% (95% CI, 81.81% to 99.34%), and the specificity was 88.89% (95% CI, 51.75% to 99.72%). The overall accuracy of the bCUBE qPCR platform for detecting leg samples was 93.48% for ZIKV (95% CI, 82.10 to 98.63%) and 86.96% for DENV2 (95% CI, 73.75% to 95.06%) (Fig. 4).

**Fig. 4.**
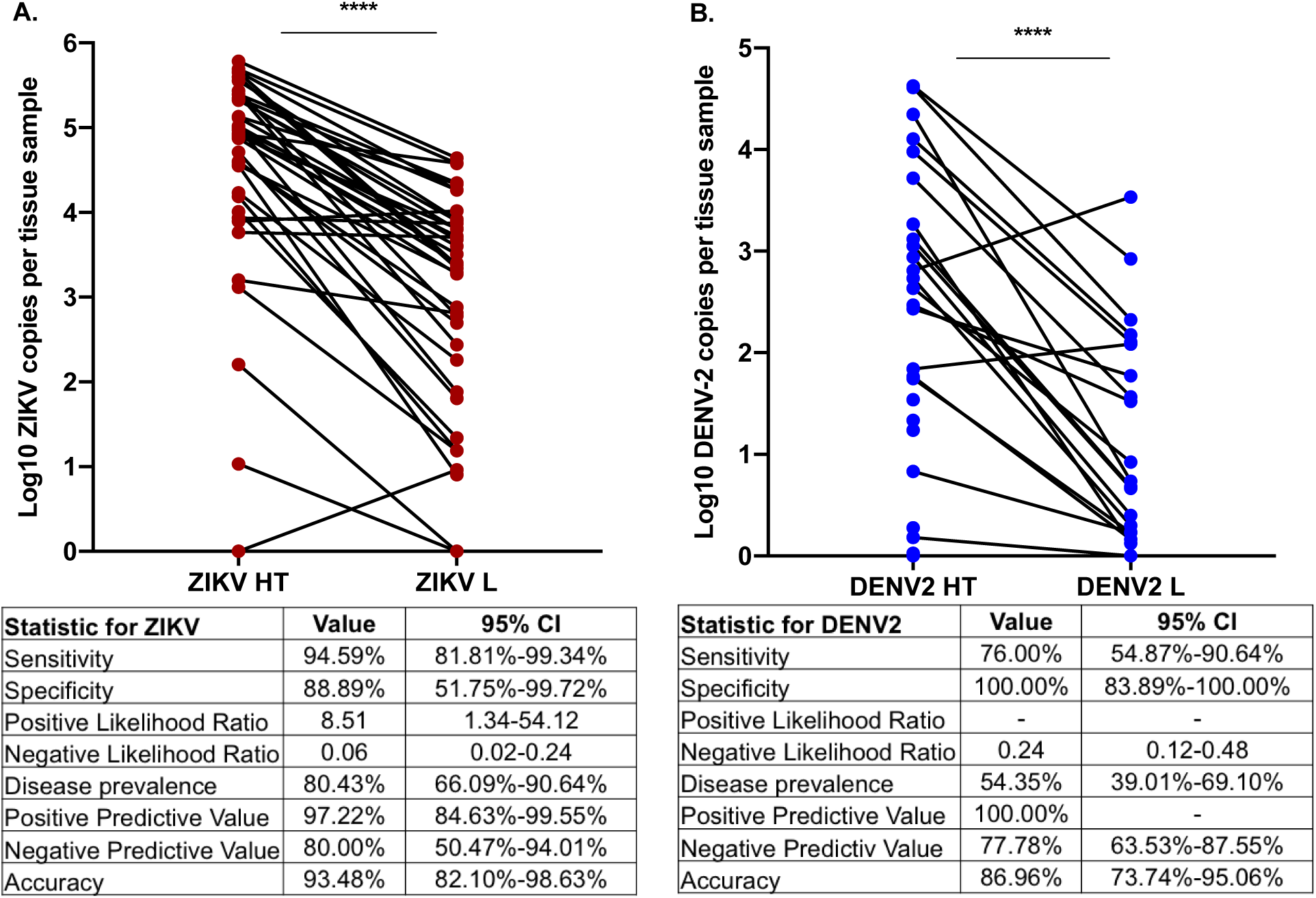
Viral concentration of individual head with thoraces compared to legs. Infected *Ae. aegypti* (n=46) were collected, and identical samples of head with thorax and legs were amplified for Zika virus (A) and DENV2 (B) using bCUBE-based qRT-PCR. Infection status was compared (****, p<0.0001; paired two-tailed t-test).

### bCUBE-based qRT-PCR detects ZIKV and DENV-2 viral RNA in pooled *Ae. aegypti* samples

Surveillance programs monitoring ZIKV and DENV2 transmission will typically assay the presence of virus in pooled samples ranging from 5 to 50 mosquitoes (50). We pooled individual ZIKV- or DENV2-infected mosquitoes with 5, 10, 15, or 20 uninfected *Aedes* in order to assess the assay’s sensitivity with pooled samples. The infection status of individual infected mosquitoes was confirmed by viral RNA extraction using squash buffer followed by bCUBE-based qRT-PCR. Following confirmation of infection, positive mosquitoes were placed into four different pools for further viral detection. ZIKV could be detected in all pooled samples. DENV2 could be detected in pools of 5, 10, and 15 mosquitoes, but DENV2 viral RNA could not be detected in one of the three biological replicates of the 20-pooled mosquitoes. We found significant differences between the individual infected samples and pooled samples; however, the detection in pooled samples was still within the range for a positive identification (Supplementary Table 1).

### bCUBE-based qRT-PCR detects *Wolbachia* in individual *Ae. aegypti*

*Wolbachia w*AlbB was amplified from individual mosquitoes with qRT-PCR using the bCUBE platform (n=35) (Supplementary Table 2). Uninfected *Ae. aegypti* (n=10) were used as negative controls. All *Wolbachia*-infected mosquitoes were positive for *Wolbachia* infection, and the negative controls did not amplify. These data suggest that bCUBE qPCR can be used for *Wolbachia* detection employing previously established primers.

## Discussion

Arboviral pathogens represent an expanding threat across the world. Monitoring the vector and its infectious cargo is essential for planning adequate mosquito control to reduce transmission. Endemic countries frequently lack surveillance resources for monitoring pathogen and vector distribution. Therefore, cost-effective field-deployable pathogen detection methods can greatly facilitate surveillance and control efforts. We sought to develop a method for detecting viral pathogens in their *Aedes aegypti* vector with the potential for field deployment. Currently, pathogen detection in the field is limited by expensive lab-based equipment and reactions and the necessity for highly trained personnel to process samples. Real-time qPCR has long been considered the gold standard for viral detection; however, inadequate access to sample preparation methods and the requirement for bulky equipment render this method incompatible with fieldwork.

As an alternative to laboratory PCR, we have now explored the use of bCUBE, a novel qPCR platform, with a simple nucleic acid extraction method for detection of arboviruses and *Wolbachia* in mosquito vectors. Actions were taken to limit the cost needed for analysis. We used DNA dye binding qPCR commercial kits as an alternative to hydrolysis probe-based qPCR kits. Furthermore, we extracted DNA/RNA using the squash buffer methodology as opposed to expensive extraction kits. It is important to note that current bCUBE cartridges are restricted to 16-wells thereby limiting the number of samples run during each experiment. Hyris bCUBE software can be programmed to display results as positive or negative result as opposed to interpreting cycle threshold and melting curve values. This minimizes training for remote personnel.

We confirmed the reliability of the bCUBE by comparing its performance against laboratory standards. Therefore, bCUBE is capable of performing real-time qPCR while overcoming the barriers presented by the need for bulky equipment. We then employed a crude extraction method using squash buffer to extract viral DENV2 and ZIKV RNA from mosquitoes for use in the bCUBE system. To address challenges involving primer cross-reactivity with other arboviruses (46, 48) as well as the tendency toward false-positive amplification in negative samples, we tested the specificity of the assay against a panel of frequently co-circulating arboviruses. The results confirmed the assay’s specificity for DENV2 and ZIKV RNA, with the potential to apply these primer pairs for field conditions. To negate any non-specific amplification of the host’s genetic material, we used a standard curve to determine the limit of detection and subsequently, the cut off values for detecting ZIKV and DENV2. This allowed for positive identification of viral RNA from infected mosquito samples while limiting the amplification of false positive samples as a result of background noise. Following primer optimization, we compared the bCUBE assay to plaque assays for determining infection prevalence. Because qPCR is generally capable of detecting low RNA copy numbers in individual samples, it is important to evaluate the assay’s potential for detecting infectious viral RNA rather than viral RNA that is not replicating (49, 51). No significant differences were seen between the bCUBE and plaque assay for three biological replicates, suggesting that the bCUBE assay is able to detect infectious viral RNA.

We evaluated the use of *Ae. aegypti* leg samples for viral detection. Some infected leg samples turned out to be false negatives suggesting this sample type is unreliable for assaying virus. However, mosquito surveillance agencies typically evaluate arboviral infection status in pooled samples as opposed to assaying individual mosquitoes. ZIKV RNA was detected in pools of 5, 10, 15, and 20 with uninfected mosquitoes in three biological replicates. DENV RNA was not detected in one of three biological replicates of 20-mosquito pools, suggesting possible limitations for DENV detection. This highlights the bCUBE platform’s potential for pool testing in the field.

Given RNA’s inherent instability and frequency of mutation, it is likely for sequence variability to occur under field conditions. Therefore, it is important to vigilantly monitor the viral strains that are co-circulating in a region to accurately and specifically detect viral RNA in mosquitoes when applying this methodology in the field. It is critical to perform absolute quantification analyses and determine end-point limitation values during optimization to avoid amplification of false positive samples as a result of qPCR background noise. Potential future studies involve bringing the portable platform into the field to analyze its feasibility for performing qPCR under true field conditions. The versatility of the assay can be applied to future studies involving *Plasmodium falciparum* detection in *Anopheles* species and monitoring of various species of the *Anopheles* vector as well as detecting insecticide resistance in field-caught mosquitoes.

## Conclusion

We have developed a highly sensitive qPCR method for detection and quantitation of DENV2 and ZIKV in single mosquitoes using the portable qPCR Hyris bCUBE platform. Our assay allowed for sensitive and reliable DENV2 and ZIKV detection that make it a potential tool for mosquito surveillance programs in endemic countries facing arboviral outbreaks. The detection of *Wolbachia* in *Ae. aegypti* further demonstrates the platform’s ability to be used as a multi-detection assay. By overcoming the challenges of costs associated with reagents and equipment, the bCUBE qPCR platform provides a potential field-deployable laboratory resources for mosquito surveillance agencies in remote regions.

## Acknowledgments

The authors thank Hyris, Inc. for providing the bCUBE and Mr Stefano Lo Priore and Mr Lorenzo Colombo for technical advices. We also thank Dr. Zhiyong Xi from Michigan State University for kindly providing *Wolbachia w*AlbB-infected *Ae. aegypti* strain WB1 and Dr. Deborah McClellan for editorial assistance. We thank the Johns Hopkins Malaria Research Institute Insectary core facility. This work has been supported by the National Institutes of Health grant R21AI136456, and the Bloomberg Philanthropies.

## Notes

### Competing Interest Statement

The authors have declared no competing interest.

### Summary of Updates

A heading for figure 4 has been added.

