## Supplemental Figure S1 for "Field-deployable molecular diagnostic platform for arbovirus and *Wolbachia* detection in *Aedes aegypti*"

**Additional File 1: Fig. S1.**

**Ct standard curves for DENV2 and ZIKV generated using GoTaq 1-Step RT-qPCR (Promega)**

Absolute quantification was based on standard curve analyses using cloned fragments from the DENV2 and ZIKV stocks. Viral RNA was adjusted to $10^{10}$ and serially diluted 10 times for qRT-PCR. Cycle threshold (Ct) values are plotted against log of RNA copy numbers (RNA copies/µl).

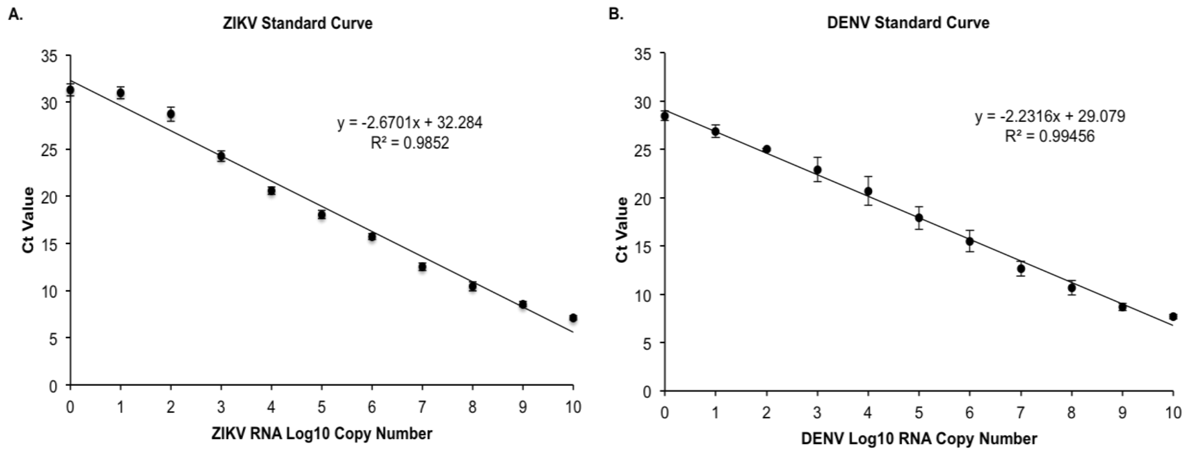
