## Supplemental Table S1 for "Field-deployable molecular diagnostic platform for arbovirus and *Wolbachia* detection in *Aedes aegypti*"

**Additional File 2: Table S1.**
**Cycle threshold values of *Wolbachia* infected *Aedes aegypti.***
Previously developed primers were used for SYBR green qPCR on the Hyris bCUBE platform to amplify *Wolbachia* infected *Aedes aegypti* (n=35). Ct values ranged from 18.92 to 26.48.

| Sample | Ct Values |
|---|---|
| W1 | 21.49 |
| W2 | 24.86 |
| W3 | 24.26 |
| W4 | 21.72 |
| W5 | 21.27 |
| W6 | 26.37 |
| W7 | 25.18 |
| W8 | 22.51 |
| W9 | 23.92 |
| W10 | 25.93 |
| W11 | 22.15 |
| W12 | 21.49 |
| W13 | 23.32 |
| W14 | 26.48 |
| W15 | 23.87 |
| W16 | 24.63 |
| W17 | 23.57 |
| W18 | 22.71 |
| W19 | 23.15 |
| W20 | 26.35 |
| W21 | 19.03 |
| W22 | 19.03 |
| W23 | 19.15 |
| W24 | 19.00 |
| W25 | 19.07 |
| W26 | 19.38 |
| W27 | 19.21 |
| W28 | 18.94 |
| W29 | 18.92 |
| W30 | 18.96 |
| W31 | 18.99 |
| W32 | 19.11 |
| W33 | 19.38 |
| W34 | 19.17 |
| W35 | 19.95 |
