## Supplemental Table S2 for "Field-deployable molecular diagnostic platform for arbovirus and *Wolbachia* detection in *Aedes aegypti*"

**Additional File 3: Table S2**

**Detection of viral RNA in pooled *Aedes aegypti* samples by bCUBE**

Pooled samples were prepared by combining individual ZIKV- and DENV2-infected mosquitoes with pools of uninfected mosquitoes. Infection of individual mosquitoes was confirmed prior to adding the individual samples to the pooled samples. Pools of 5, 10, 15, or 20 uninfected mosquitoes plus one infected mosquito were evaluated by bCUBE. Table lists RNA concentration of three biological replicates of pooled samples with corresponding concentration of the individual mosquito added.

| ZIKV Pooled Number | RNA $log_{10}$ Pooled | RNA $log_{10}$ Corresponding Individual |
|---|---|---|
| **5** | 3.22 | 4.26 |
| **10** | 3.26 | 5.54 |
| **15** | 3.62 | 5.25 |
| **20** | 3.99 | 5.55 |
| **5** | 1.91 | 3.59 |
| **10** | 1.80 | 4.20 |
| **15** | 2.35 | 3.99 |
| **20** | 2.77 | 4.40 |
| **5** | 1.15 | 1.64 |
| **10** | 0.90 | 1.68 |
| **15** | 1.62 | 3.79 |
| **20** | 1.64 | 3.44 |

| DENV2 Pooled Number | RNA $log_{10}$ Pooled | RNA $log_{10}$ Corresponding Individual |
|---|---|---|
| **5** | 3.23 | 5.18 |
| **10** | 1.16 | 4.32 |
| **15** | 2.20 | 4.44 |
| **20** | 2.45 | 4.77 |
| **5** | 1.24 | 3.17 |
| **10** | 1.68 | 2.79 |
| **15** | 0.51 | 3.27 |
| **20** | 1.73 | 4.11 |
| **5** | 1.54 | 2.88 |
| **10** | 0.26 | 1.69 |
| **15** | 0.16 | 1.95 |
| **20** | -0.33 | 1.67 |
